# Paying Attention to Attention: High Attention Sites as Indicators of Protein Family and Function in Language Models

**DOI:** 10.1101/2024.12.13.628435

**Authors:** Gowri Nayar, Alp Tartici, Russ B. Altman

**Affiliations:** Department of Biomedical Data Science, Stanford University; Department of Genetics, Stanford University; Department of Medicine, Stanford University; Department of Bioengineering, Stanford University

## Abstract

Protein Language Models (PLMs) use transformer architectures to capture patterns within protein sequences, providing a powerful computational representation of the protein sequence [1]. Through large-scale training on protein sequence data, PLMs generate vector representations that encapsulate the biochemical and structural properties of proteins [2]. At the core of PLMs is the attention mechanism, which facilitates the capture of long-range dependencies by computing pairwise importance scores across residues, thereby highlighting regions of biological interaction within the sequence [3]. The attention matrices offer an untapped opportunity to uncover specific biological properties of proteins, particularly their functions. In this work, we introduce a novel approach, using the Evolutionary Scale Model (ESM) [4], for identifying High Attention (HA) sites within protein sequences, corresponding to key residues that define protein families. By examining attention patterns across multiple layers, we pinpoint residues that contribute most to family classification and function prediction. Our contributions are as follows: (1) we propose a method for identifying HA sites at critical residues from the middle layers of the PLM; (2) we demonstrate that these HA sites provide interpretable links to biological functions; and (3) we show that HA sites improve active site predictions for functions of unannotated proteins. We make available the HA sites for the human proteome. This work offers a broadly applicable approach to protein classification and functional annotation and provides a biological interpretation of the PLM’s representation.

**Author Summary:** Understanding how proteins work is critical to advancements in biology and medicine, and protein language models (PLMs) facilitate studying protein sequences at scale. These models identify patterns within protein sequences by focusing on key regions of the sequence that are important to distinguish the protein. Our work focuses on the Evolutionary Scale Model (ESM), a state-of-the-art PLM, and we analyze the model’s internal attention mechanism to identify the significant residues.

We developed a new method to identify “High Attention (HA)” sites—specific parts of a protein sequence that are essential for classifying proteins into families and predicting their functions. By analyzing how the model prioritizes certain regions of protein sequences, we discovered that these HA sites often correspond to residues critical for biological activity, such as active sites where chemical reactions occur. Our approach helps interpret how PLMs understand protein data and enhances predictions for proteins whose functions are still unknown. As part of this work, we provide HA-site information for the entire human proteome, offering researchers a resource to further study the potential functional relevance of these residues.

## 2 Introduction

Understanding a protein’s characteristics from the sequence is crucial to predicting its function, and analyzing protein data at scale requires computational representations of the protein sequence [5, 6]. Protein Language Models (PLMs) are computational tools that apply transformer-based architectures, originally developed for natural language processing, to protein biology [1]. These models are trained on large datasets of protein sequences to learn meaningful representations of protein sequences by capturing patterns across the amino acid sequence. A product of the PLM is a numerical representation of the sequences called embeddings. The embeddings encapsulate information about the proteins’ biochemical and structural properties, making PLMs valuable for a variety of downstream tasks, such as predicting protein function, classifying proteins into families, and exploring structural relationships among proteins [7, 8, 9, 10]. Within the PLM model, the attention mechanism is the key innovation that facilitates capturing longrange dependencies and contextual relationships across the sequence [11]. The attention mechanism uses Query (Q), Key (K), and Value (V) matrices to compute attention scores, which determine the importance of each amino acid by weighing its interactions with other residues in the sequence [3].

The PLM utilizes multiple layers of attention, and the model refines its understanding of the sequence over the layers, improving the depth and accuracy of the learned representations. Attention matrices produced at each layer of the model reflect how the PLM interprets the importance of specific residues within a protein sequence [12]. Therefore, the attention mechanism within PLMs, which assigns weights to each amino acid (AA) in a sequence based on its relevance to others, contains crucial information about which residues may be important to the proteins’ function and structure [13, 14]. However, previous work has not fully explored how these attention patterns correlate with known biological functions or the classification of proteins into families [5, 15].

Proteins with similar sequences tend to have similar representation vectors in the high-dimensional space generated by PLMs [15, 16], which suggests that the model recognizes common features among these proteins. Therefore, within the attention layers, the model must converge to identify the similarities in the sequence [17]. Thus, we hypothesize that investigating the parts of the sequence that the model prioritizes will provide insights into the key residues that are important to the protein’s biological properties.

In this work, we develop a method to systematically identify key residues—termed High Attention (HA) sites and show that these sites are indicators of the protein family and function, described graphically in Figure 1. We specifically focus on a state-of-art PLM, Evolutionary Scale Model 2 (ESM-2) [4], which was trained on a corpus of protein sequences and provides the attention values at each layer, embedding vector for each residue in the sequence, and the predicted structure contact map for the sequence [13]. Our approach identifies the specific layer in the PLM where attention converges on these HA sites, providing a clearer understanding of how the model distinguishes between different protein families. By focusing on these key residues, we demonstrate that these HA sites can be used to better predict protein function and classify proteins into families. We also enhance the interpretability of PLMs, by identifying the biological significance of the computational representations within the model. The major contributions of this work are:

1. Identification of High Attention Sites (HA sites): We develop a robust method to identify the specific residues in protein sequences that receive the most attention in the middle layers of the PLM, which are critical for the model’s family classification decisions.
2. Protein Family Identifier: We show how the attention patterns in PLMs can be analyzed to provide a clearer understanding of the model’s predictions, linking HA sites to distinguishing features of protein families, specifically biological functions and protein sequence and structure.
3. Enhanced Functional Prediction: We demonstrate that using HA sites improves the prediction of protein functions, particularly for proteins with previously unknown functions, offering a dataset of predicted, functionally important residue annotations across the human proteome.
4. We make the set of HA sites available for the entire human protein, along with the alignment based on the HA sites for all human protein families, to facilitate studying key sites in each protein family.

**Figure 1:**
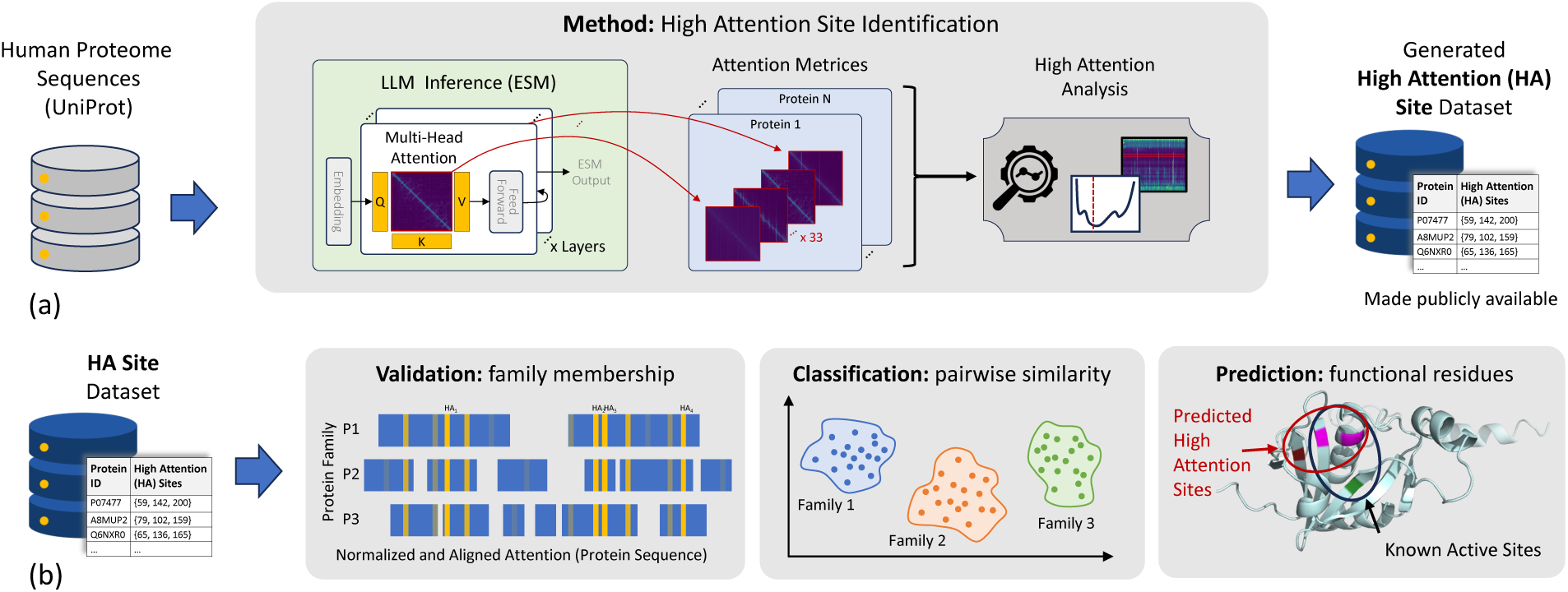
Overview diagram of the method and contributions of this work. Panel A shows a high-level description of the data and method used to identify high-attention (HA) sites. We use the ESM model to analyze all human proteins, by running the protein language model on all human protein sequences and obtaining the attention matrices. We implement our attention analysis algorithm (described in Section 3.1) to obtain a set of HA sites for each protein. This dataset of HA sites is made publicly available. Panel B shows three tasks that can be performed with the set of HA sites. (1)The HA sites can be used to confirm the protein family alignment and validate a protein’s membership within a family based on the similarity of the progression of attention values. (2)The HA sites can define vectors that can be used to determine pairwise similarities between proteins. (3)The HA sites are also predictive of functionally important regions of the protein, as they are spatially close to active sites and residues crucial to the specificity of the protein.

## 3 Background

### 3.1 Protein Language Models

Large language models for proteins (PLM) draw from the transformer-based advancements seen in natural language processing and are trained on the universe of protein sequences to learn a numerical representation for the amino acid sequences. These numerical representations for proteins can be used to predict the properties of protein sequences [5]. PLMs are trained on the protein sequence to capture complex patterns within the amino acid (AA) sequences, and the sequence is passed through layers of attention that score the importance of a given token to the others within the sequence. Each AA is treated as a distinct token within the transformer network, and thus the PLM is able to learn the contextual relationship between AAs. The PLM generates an embedding, or vector representation, for each of the AAs within a given protein sequence that can be used for downstream prediction tasks, such as structure prediction [9, 10].

### 3.2 Representation Vectors

PLMs generate a vector for each token (amino acid) in an input protein sequence, encapsulating the contextual information derived from the sequence. These token-level vectors capture biochemical and structural properties within the sequence context [18]. To obtain a single representation vector for the entire protein, various pooling strategies are employed, such as mean pooling, max pooling, or using a special classification token (CLS). Mean or max pooling aggregates token vectors by either averaging their values or selecting the maximum value across all tokens. The CLS token is an artificial token introduced in training that is updated based on every other token, thus aggregating information across the sequence into a summary vector [19].

These pooled representation vectors have been used to identify similarities between proteins, allowing for the classification of proteins into families and the prediction of their functions [19, 14]. However, these pooling techniques consider each AA token to be of equal importance to the protein function, which is biologically incorrect AAs at specific residues within a protein sequence can have a much higher impact on the overall structure and function than others [20, 12]. Instead, the attention matrix, from which the representation vector is derived, provides residue-level information. By preserving this granular data, we can identify specific sites within the protein that are playing functionally important roles.

### 3.3 Attention Matrices

Attention mechanisms dynamically weigh the importance of different tokens (amino acids) relative to each other. Attention matrices are computed by first generating Query (Q), Key (K), and Value (V) matrices from the input amino acid sequence through learned linear transformations [3]. The attention scores are then calculated by taking the dot product between the Query and Key matrices, which are subsequently normalized (typically with softmax) and used to weigh the Value matrix, producing the final attention output that highlights the most contextually relevant amino acids in the sequence. PLMs typically have multiple layers of attention to progressively refine the sequence representation [17]. Each layer within a PLM generates an attention matrix, where each matrix entry reflects the significance of one amino acid’s influence on the representation of another. The last layer of attention is used to generate the token representations, and each layer of attention uses the attention from the prior layer [21]. These matrices allow the model to capture long-range dependencies within the protein sequence, creating a context-aware representation of the protein. Thus, the PLM builds a complex model of the dependencies between positions in the sequence with the attention matrices, allowing downstream predictions of other protein features that are sequence-dependent[1]. The attention matrices provide residue-level importance at each layer, making them invaluable for identifying critical functional residues [22, 23]. This insight leads to the hypothesis that attention matrices can be leveraged to better identify functionally important residues within a protein.

Importantly, the representation vector for each protein is derived from the layers of attention within the model, with each layer contributing to the final embedding. The representation vector for proteins with similar sequences cluster together in the high-dimensional space [15, 9]. Since the representation vector is a result of the attention matrices, this implies that the attention matrices for similar protein sequences converge as the sequences progress through the transformer layers. In this work, we show that this convergence suggests that the model consistently identifies similar key residues across similar protein sequences, and highlight the relationship between the key residues and the function of the protein.

### 3.4 Protein Families

Protein families are groups of proteins that share a common evolutionary origin, typically reflected in similarities in their amino acid sequences, structures, and often, their biological functions [24]. Members of a protein family are usually characterized by conserved regions, which are critical for maintaining the protein’s function. These conserved regions often correspond to active sites, binding sites, or structural motifs that are essential for the protein’s biological activity. The protein families have been annotated and are stored in publicly available databases, such as PFAM [25].

Traditional methods for classifying proteins into families rely on sequence alignment techniques, such as multiple sequence alignment (MSA), which identify consensus sequences across different proteins [26]. These consensus sequences often indicate the regions of the protein that are important to the function or structure of the family [20]. Structural similarity is measured by evaluating the distances between residues in the three-dimensional structure, which is stored in the Protein Data Bank (PDB) [27, 4]. The functional similarity is often computationally measured by comparing known functional annotations; the Gene Ontology (GO) is one database that provides functional annotation codes for each protein [28, 29]. The convergence of the attention matrices can also be used as a signal for similar proteins, and so in this work we evaluate the significance of similar regions of the attention matrices on the characteristics that define a protein family function, sequence, and structure.

### 3.5 Evolutionary Scale Model

In this work, we specifically focus on the Evolutionary Scale Model (ESM), a state-of-the-art PLM. We specifically use ESM version 2 with 650 parameters, which uses 33 layers of attention [13, 4]. ESM is a bidirectional transformer, which calculates attention by simultaneously considering context from both preceding and succeeding tokens in a sequence, enabling it to capture dependencies in both directions for more comprehensive representation learning [21]. ESM2 also uses 14 attention heads in each layer; each attention head has its own set of Q, K, and V matrices, which are independently learned for each attention head. The use of attention heads allows the model to capture different aspects of the sequence simultaneously, as each head uses different learned linear transformations.

We use ESM outputs in this study because ESM provides access to the representation vectors for each token, the attention matrix for each layer, and the predicted contact map for each input sequence. A protein contact map represents the distance between all possible amino acid residue pairs in a protein structure, and it is predicted directly from the attention matrices, creating a proxy for the predicted structure of a sequence. The ESM model generates the predicted contact map by performing regression over the layers of attention matrices [13, 30]. We study the ESM model because of the ability to test the relationship between the attention matrices and the structure through the generated contact maps, which are not provided by other PLMs. However, given that other bidirectional transformer PLMs follow a similar structure of layers of attention, the concept of evaluating the similarities in attention matrices across the layers to find similarly functioning proteins can be generalized across models[31].

## 4 Method

We identify key residues that influence PLM representations of similar proteins. Our method systematically detects the earliest PLM layer with consistent attention matrices across a protein family, using it to pinpoint high-attention (HA) sites. We then assess how well the HA sites capture the family’s defining characteristics: function, sequence, and structure.

### 4.1 Identifying the convergence layer and key residues from attention matrices

We use the ESM-2 650-million parameter model on all human protein sequences obtained from UniProt [32]. We run ESM-2 and obtain the attention matrix for each layer (L = 33 layers) for each attention head (H = 14 attention heads) for each protein. Because each attention head uses different learned linear transformations, the attention heads are attuned to different aspects of the sequence. Since this work focuses on analyzing the change in attention as the sequence progresses through the model (through the layers) rather than across heads, we first pool the attention heads (H) to create one attention matrix per layer. We perform mean pooling, taking the mean across all the attention heads at each (*i, j*) cell in the matrices, where *i, j* are two residue sites in protein *p*. This pooling ensures that the analysis reflects the overall attention distribution at each layer rather than head-specific biases.

The columns of the attention matrix denote the importance of the token AA at a given column on all other tokens in the sequence. Thus, to quantify the global importance of one residue, we perform a column-wise summation of the pooled attention matrix. We create a vector, *v_l_* with values in the range 0 to 1 for each attention layer, such that the value in position *i* in the vector, *v_l_* denotes the relative importance of the token AA at position *i* in the sequence in layer *l*. We normalize this vector to values between 0 and 1 to allow for comparison between the residues, as we aim to identify the residues with the highest relative attention. The operations are described in equations 1 − 3 and in Figure 2.

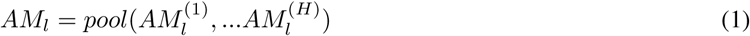

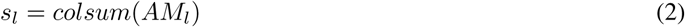

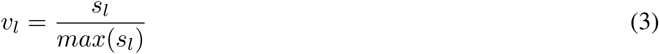

**Figure 2:**
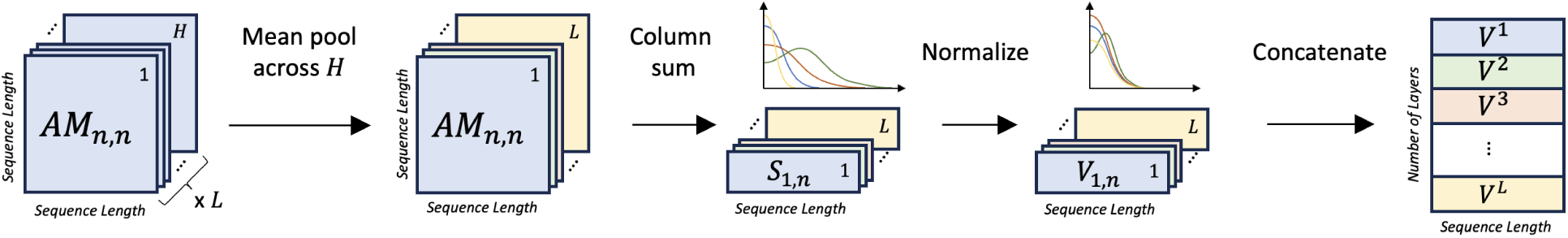
Schematic diagram of the calculations performed on the attention matrices to create the normalized vector per layer. For one protein of length *n*, all attention matrices are obtained from the PLM and we first mean pool the attention heads, *H*, for each layer, *L*. This leads to a *n* × *n* matrix for each layer. We do a column sum of this matrix to measure the importance of each residue across the sequence and create a 1 × *n* vector for each layer. We normalize this vector to facilitate the comparison of the attention values across proteins. We then concatenate these per-layer vectors into a matrix of size *L* × *n*, to evaluate the change in attention for each residue over the layers.

HA sites are defined as the residues with normalized attention values approaching 1, while the normalized attention values for all other residues in the sequence approach 0. We call the first layer manifesting this bimodal attention distribution the convergence layer and it is the first layer that the PLM identifies the category, or protein family, to which this sequence belongs. To systematically identify the convergence layer, we sort the values of *v_l_* in descending order for each layer *l*. To satisfy the HA site constraints such that there is the largest delta in attention between the highattention residue and the other resides, we want to identify the layer at which the sorted values have the sharpest point of inflection. To do this, we fit two lines to the sorted *v_l_*, one for the initial, higher attention residues and another for the remaining lower attention residues. The angle *θ* between these two lines is calculated for each layer. The layer with *θ* closest to 90 degrees has the largest separation between high and low attention residues, indicating the convergence layer and the emergence of HA sites. The indices of these high-attention residues the indices corresponding to the values to the left of the point of intersection of the two fit lines correspond to the HA sites.

### 4.2 Using HA sites to measure similarity of proteins

We show that the HA sites can characterize a protein family and thus can be used to determine a similarity relationship between two proteins. We develop this method to enable comparing proteins of varying lengths while maintaining residue-level granularity.

To compare two proteins, we compare the normalized attention of their HA sites over the layers of the PLM. For each HA site in a protein, we create vector *t*, such that the *i^th^*index of *t* is the normalized, summed attention value (described in Section 3.1) of the given HA site residue at layer *i*. For a given protein with HA site at residue *m*, *t*[*i*] = *v_i_*[*m*]. We generate one such vector for each HA site in a protein and to compare two proteins, we calculate the pairwise cosine similarity between the two sets of HA sites. We then take an average of the similarities to create a similarity metric for the two proteins. With this method, the *t* vectors are the dimension of the number of layers of the PLM (L), and thus, this method does not depend on the length of the protein.

We evaluate the ability to distinguish between proteins in a known family against proteins randomly chosen from the set. We calculate the distribution of scores from within the same family (in-family) and compare it to the distribution of scores between randomly chosen proteins (out-family) and perform a Kolmogorov–Smirnov (KS) test to determine the significance of the difference between the distributions. We also evaluate the impact of the specific HA sites within one family by comparing the scores resulting from the true HA sites to the scores when using random sites as overlap. This allows us to evaluate if the true score is significant in comparison to a background distribution. We perform the same KS comparison using the cosine distance between the mean-pooled representation vector, max-pooled representation vector, and CLS-token vectors (described in Section 2.2)

### 4.3 Evaluate the relationship between HA sites and active sites

Protein families are a useful functional model for organizing similar proteins; proteins that share sequence or structure typically share function.

The active site of a protein is crucial to the protein’s function as it is the region where substrate molecules bind and undergo a chemical reaction. Therefore, we evaluate the correlation between the active sites and HA sites of a protein.

From UniProt, we obtain the active site residues. Using the protein structures obtained from the Protein Data Bank, we compare the distance in three-dimensional space between the active site and HA site. We use the coordinate of the CA atom for the HA-site residue and the active site residue and calculate the euclidean distance between these two coordinates. If a protein has multiple HA sites and active sites, the closest pair is considered. For the protein families with no active site annotation in UniProt, we perform a qualitative analysis of the HA sites as a prediction of their active site.

### 4.4 Evaluating HA sites correlation to known definitions of protein families (sequence and structure)

Traditionally, the sequence and structure of proteins are used to define a protein family, and so we compare the HA sites to the sequence and structure motifs that classically define a protein family. We perform a multiple sequence alignment using ClustalOmegaCommandline [33] package from BioPython, which uses a progressive alignment strategy for each protein family. We find the percent consensus of residue identity and compare the consensus at HA sites to the average consensus across the alignment.

To evaluate the correlation between the HA sites with the protein structure, we use the contact map generated from ESM to compare the correlation between the attention matrix values and the contact map. The attention matrix indicates the importance of one token AA on another, while the contact map indicates the physical distance between two token AAs. We calculate the spearman correlation between each (*i, j*), for all *I* ∈ *set*(HA site), in the attention matrix and the contact map, for all layers of attention to evaluate the importance of these sites on the predicted structure.

## 5 Results

### 5.1 Dataset

We download all human proteins (protein name and sequence) from UniProt, producing 20,085 proteins. From Hugging Face, we access the ESM-2 model, specifically the pre-trained model esm2 t33 650M UR50D with 33 layers, 14 attention heads per layer, and 650M parameters. We use the pre-trained model to obtain each attention matrix (14 x 33 matrices of size sequence length x sequence length), the representation vector of length 1280 for each token AA (1280 x sequence length), and the contact map (sequence length x sequence length). For each protein, we get the PFam ID, leading to 7,289 unique families. We also obtain the structure file for each protein from the Protein Data Bank (PDB), which is the largest protein structure database. When an experimentally generated structure for a protein was not present in the PDB, we used the AlphaFold2-generated structure, also stored in the PDB. Thus, we obtain the structure for 20,074 proteins. We also download the protein family annotation from Pfam for each of the human proteins.

### 5.2 Layer of convergence and HA sites are consistently identified in the middle layers of the PLM for all human proteins

Once the attention matrices for all protein sequences are generated and normalized, we evaluate the progression of attention over each token AA through the layers. We perform column normalization to evaluate the significance of a token AA to the other tokens in the sequence. We plot the resulting vector of attention values for each layer for one protein as a heatmap (rows are the layers and columns are the token AAs), where the brighter colors indicate high attention and the darker colors indicate low attention. In Figure 3, we show selected attention matrices and normalized heatmap for one example protein, P07477. Panel A shows the attention matrices from the ESM model for layers 0(first), 10, 20, and 32(last). In layer 0, all residues have similar levels of importance, for layer 10, selfattention and several high attention residues emerge. Layers 20 and 32 have a similar pattern of self-attention and show more localized attention specific to residues that are close together. Panel B shows the heatmap for P07477, after our normalization method, where each row shows the vector *v_l_* for *l* ∈ [0, 32]. The heatmap highlights the pattern seen in the attention matrices, as the early layers of the model have relatively similar levels of attention across all residues, as the normalized values for the row are all close to one. In the middle layers of the model, the the relative attention pattern changes to high attention (close to 1) for certain residues, and close to 0 attention for all other residues. The attention pattern then builds toward locally-specific attention distributed evenly across the sequence by layer 32 (last layer). The heatmaps for all proteins are included in the supplement.

**Figure 3:**
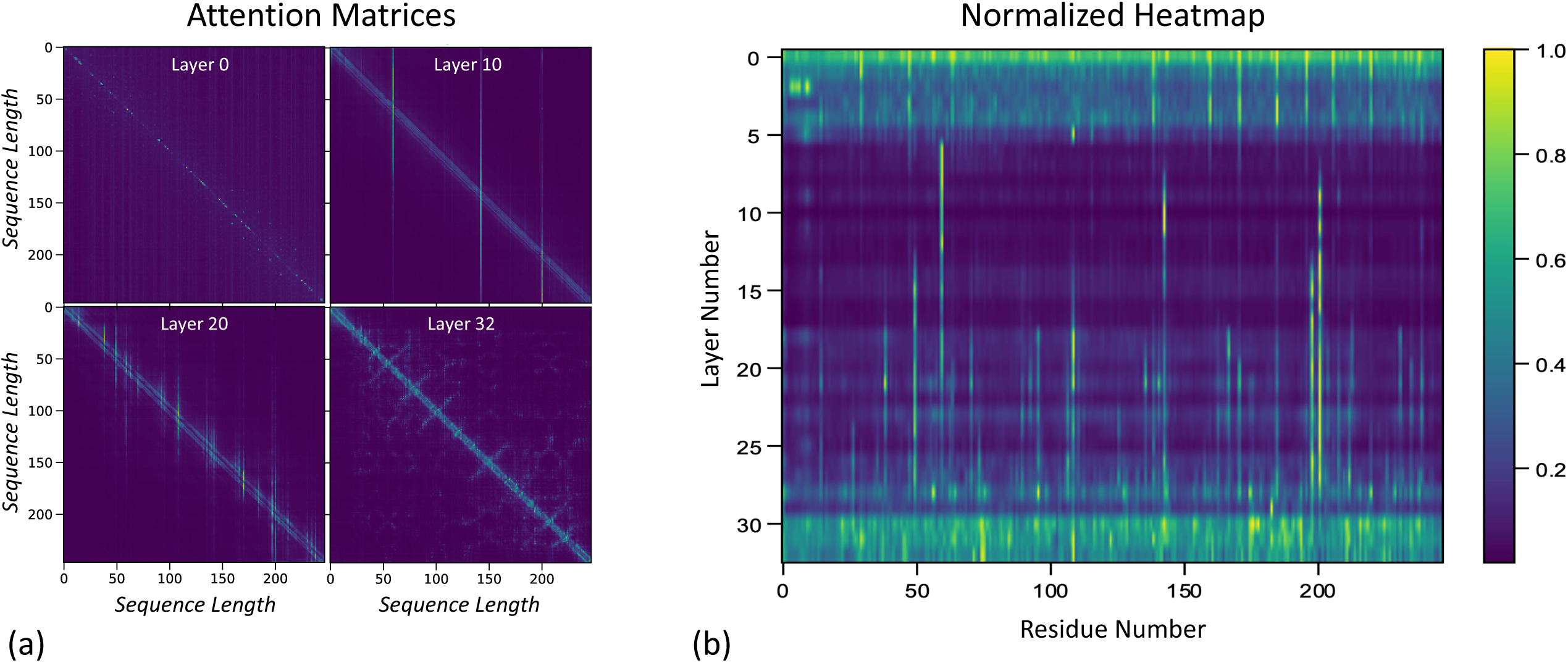
Selected attention matrices and normalized heatmap for the protein P07477. (A) Attention matrices from the ESM model at layers 0 (first), 10, 20, and 32 (last). In layer 0, all residues exhibit similar levels of importance; in layer 10, self-attention and several high-attention residues emerge. Layers 20 displays localized attention patterns, focusing on residues in close proximity. Layer 32 shows pairwise attention between residues that are close in the sequence, as well the structurally related, resembling a contact map. (B) Normalized heatmap for P07477, where each row represents the vector *v_l_* for *l* ∈ [0, 32]. The heatmap reveals attention patterns consistent with the matrices: the first layers display uniform attention across residues, while the middle layers highlight distinct residues with high attention. By layer 32, attention becomes locally specific across the sequence.

Proteins within a protein family share similar attention patterns as they progress through the network, which is made evident by comparing the normalized heatmaps between protein family members. Figure 4 show the aligned heatmaps for all members of one example protein family, Trypsin Serine Protease (PF00089).The aligned heatmaps illustrate the attention values across 33 layers for each protein, with residues aligned by their index positions. Each row corresponds to a different protein identified by its UniProt ID, and high attention values are indicated by bright colors. We show the aligned heatmaps to facilitate comparison between residues that are similar across the proteins; the empty sections of the heatmap correspond to gaps in the alignment. We observe a remarkable consistency in the attention patterns across all selected proteins. Notably, the residues with high attention follow nearly identical patterns across all proteins in the family. This consistent attention distribution suggests that the model is capturing conserved regions that are critical to the family definition, thus these regions likely correspond to sites essential for the protease activity.

**Figure 4:**
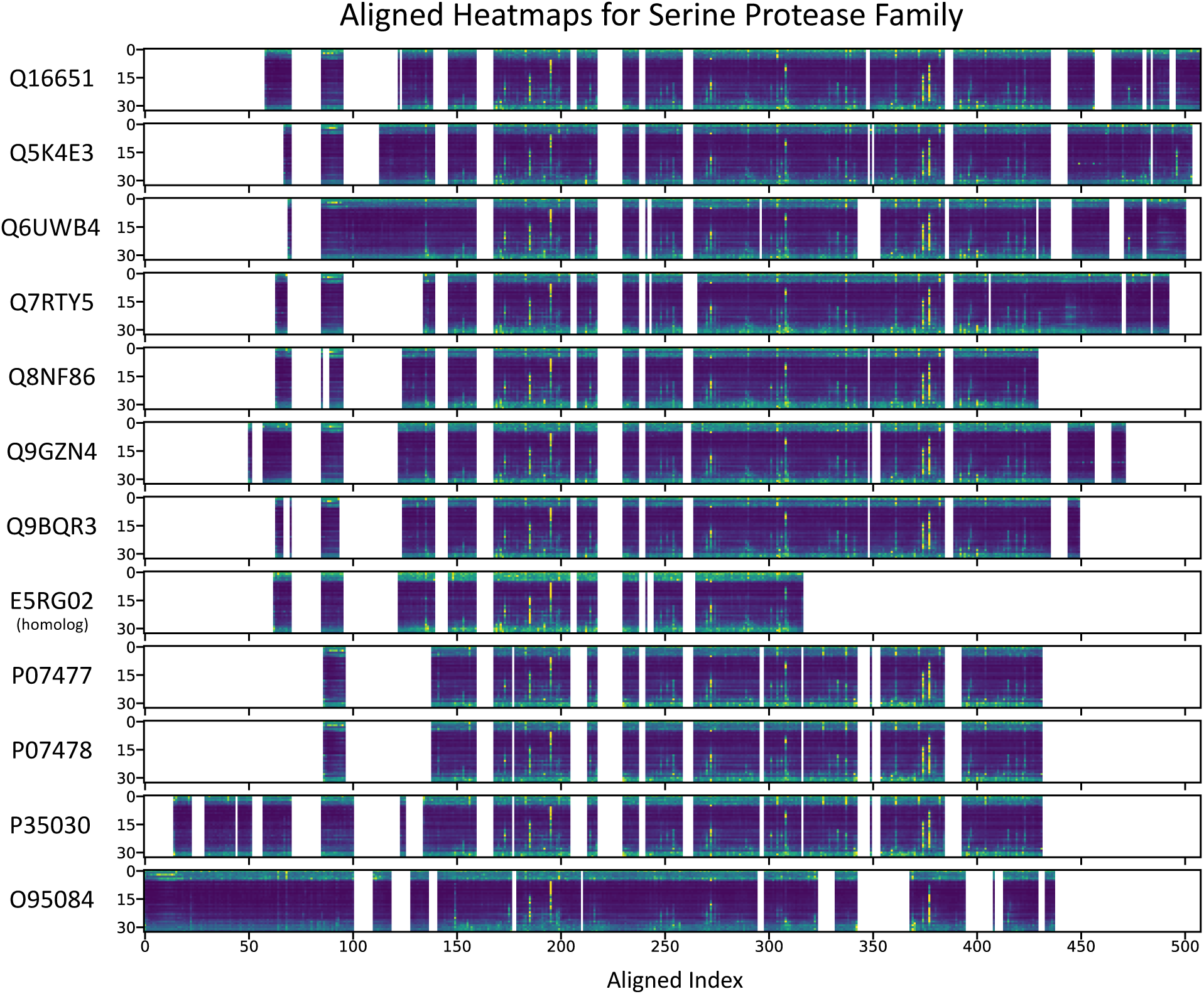
Aligned heatmaps showing the consistency of attention patterns across 33 layers for all proteins in the trypsin serine protease family (PF00089). Each row represents a distinct protein, identified by its UniProt ID, with attention values aligned by residue index. White regions indicate gaps in the alignment, dark blue regions correspond to relatively low attention, and bright yellow represents residues with high attention. Residues with high attention consistently follow the same pattern across layers for all proteins, highlighting conserved regions within the family. This strong similarity in attention distribution suggests a shared functional relevance among these high-attention residues across the family.

White regions in the heatmaps represent gaps in the sequence alignment, yet even with these gaps, the high-attention patterns remain aligned. This further emphasizes the conservation of these important residues despite minor variations in individual sequences. The protein derived from homology, E5RG02, is specifically labeled and is truncated compared to the other family members, and yet follows similar attention patterns, reinforcing the observation that the attention patterns are maintained across homologous sequences.

Figure 5 plots the ordered attention values in *v_L_* for each layer, and we show these plots for a randomly chosen protein, P70477. This figure illustrates how this method captures the shift in attention patterns across layers and identifies the HA residues. Panel (A) plots the sorted normalized attention values across each layer and the two fit lines. Here, we observe the slope of the initial attention decay, represented by the angle *θ* which varies with layer depth. In panel (B), we plot *θ* across all layers, we identify the “layer of convergence”—the layer at which *θ* is closest to 90 degrees, representing the layer with the sharpest change in values between high-attention residues and all others. For P07477, this point occurs at layer 10. Within the layer of convergence, a more detailed examination in panel (C) shows the breakpoint between the two fit lines and, thus, between high-attention and low-attention residues. This breakpoint allows us to define HA sites, which correspond to residues where attention is most concentrated. In panel (D), the layer of convergence is highlighted, with the identified HA sites shown in bright yellow. These HA sites vary from protein to protein, but the approach ensures that each site is selected based on the model’s attention distribution specific to that protein’s sequence features. This method demonstrates the model’s ability to adaptively focus on residues that are likely to be functionally or structurally significant across diverse protein sequences within the same family, providing insight into shared and unique aspects of protein function.

**Figure 5:**
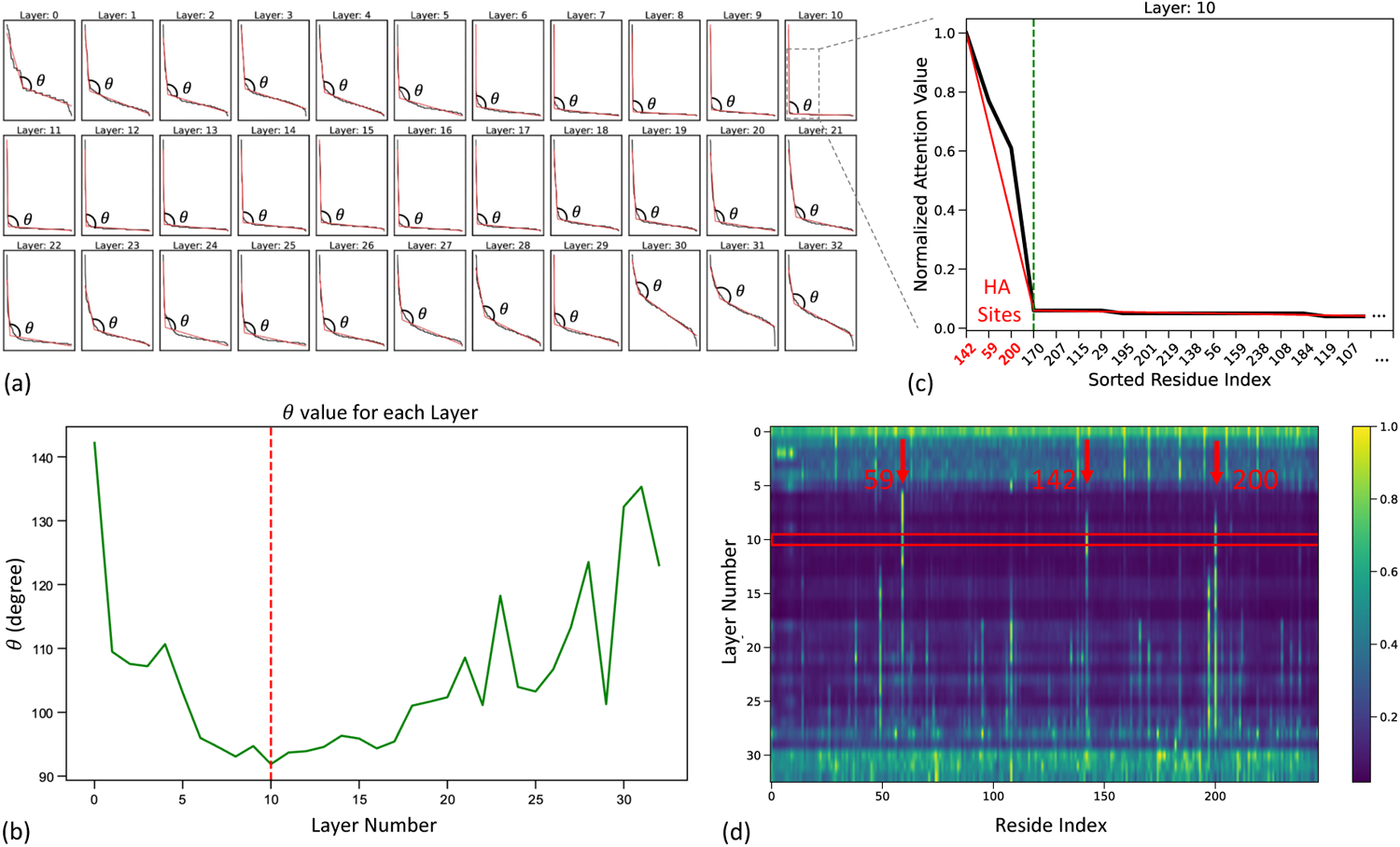
For a selected protein, P07477, an illustration of the method to identify the layer of convergence and the high attention (HA) sites. Panel (A) plots the sorted normalized attention values, *sort*(*v_l_*), in black and the two fit lines in red, and this plot is shown for each layer. Each plot shows *θ*, the angle representing the slope of the initial attention decay, which varies by layer. Panel (B) plots *θ* over all the layers in green and shows the layer with *θ* closest to 90 degree in red. This layer is defined as the layer of convergence. Panel (C) shows a zoomed-in version of the normalized attention values and fit lines for layer 10, chosen in panel B. The x-axis is truncated for readability, as the slope for the rest of the indices is close to 0, and we focus on the breakpoint between the two lines. The green dotted line denotes the break point between the two fit lines, and we choose the residues to the left of the breakpoint as the HA sites. The zoomed-in panel shows that this method identifies the breakpoint at the first low-attention residue(0.06 attention for this protein), and thus the residues to the right of this point correspond to the brightly colored columns in the heatmap. Panel (D) highlights the layer of convergence (layer 10 for this protein) with the red box and the brightly colored residues within this box (at index 59, 142, and 200) are the defined HA sites.

The number of HA sites across the human proteome ranges from 0 to 39. We find that only 1.38% of proteins do not have an identified HA site. Ninety percent of proteins have between 1 and 9 HA sites, with 50% of proteins having 1-5 HA sites. Over 50% of proteins converge to find these HA sites at layer 10. 90% of proteins converge between layers 6-14. We show a histogram of these results in Figure 6

**Figure 6:**
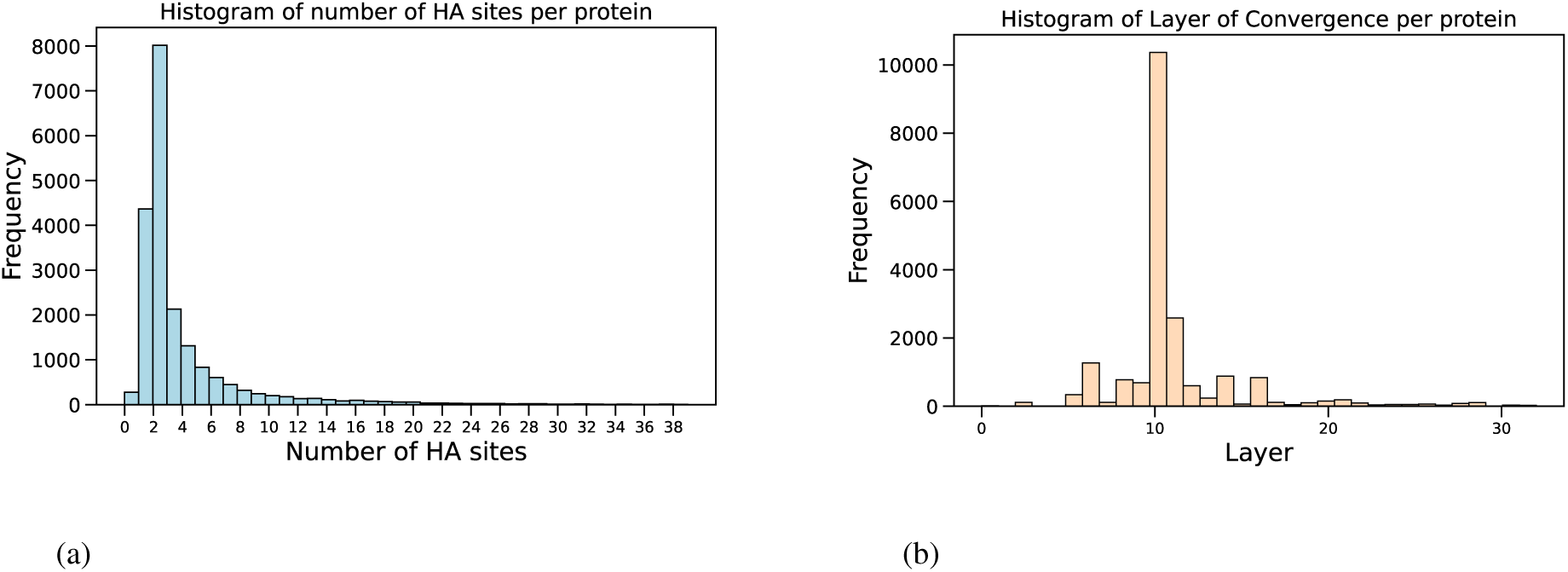
Results for HA sites and layer of convergence for all human proteins with an HA site. Panel (A) shows a histogram of the number of HA sites per protein and Panel (B) shows a histogram of the layer of convergence for each protein. The majority of proteins have 2 HA sites which emerge at layer 10, showing that for almost all proteins, the PLM converges on a set of important residues in early layers.

### 5.3 Similarity metric defined by the HA sites creates tighter clusters than pooled representation vectors

We evaluate the ability of the attention-based similarity metric (described in Section 3.2) to distinguish between inter- and intra-family proteins. We first evaluate the distribution of similarity scores created between inter and intra-family proteins using our attention-based similarity metric and compare to distributions from a cosine distance of the pooled vectors (CLS, mean, max). We show these distributions for one protein family, trypsin-like serine protease (PF00089), in Figure 7. The pairwise distances from within the family are more diverse for the CLS token and the mean-pooled representation vector. The max-pooled representation vector has a smaller range of pairwise distances within the family, but the mean within the family overlaps with the mean in the background distribution. Our HA-based similarity metric creates a small range of distributions within a family while also being separable from the random distribution.

**Figure 7:**
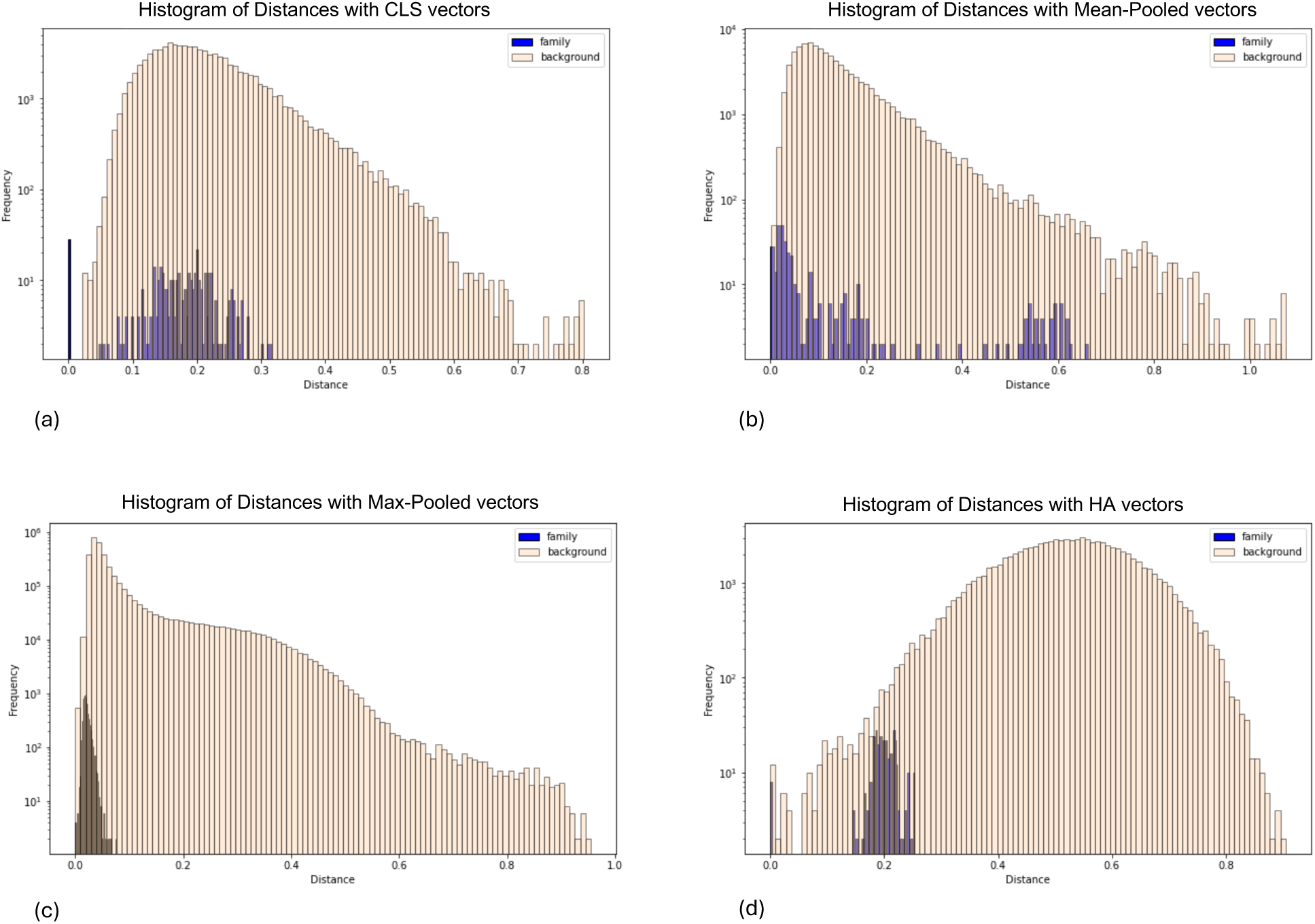
This figure shows the pairwise vector distances for one protein family, PF00089, (blue) compared to randomly chosen proteins (orange) using various similarity metrics to evaluate the utility in each similarity measure in identifying proteins from the same family. An ideal similarity metric would create an in-family distribution that is separable from the random distribution with a narrow range of distances for the in-family proteins (as we expect them to be similar to one another). For Panel A, we use the CLS vector (described in Section 2.2) for each protein and calculate the cosine distance between the two CLS vectors to determine the similarity of the two corresponding proteins. For Panel B and C, we use the mean-pooled vector and the max-pooled vector (described in Section 2.2), respectively, to define the protein and measure the similarity between two proteins by taking the cosine distance between their pooled vectors. For Panel D, we use HA-based method (described in Section 3.2) to calculate the distances between in-family and random proteins. We display the results for one protein family, serine protease. This figure shows that the in-family distribution is the tightest (smallest range of distance values) for the max pooled (Panel C) and HA-based metrics (Panel D). However, the in-family distribution for the max-pooled vectors (Panel C) is not easily separable from the background distribution, showing that even randomly chosen proteins share a low distance value with this measure. The HA-based method creates a distribution for the in-family distances that are easily separable from the background distribution, satisfying the requirements for an optimal similarity metric for family definition.

To compare the distributions, we performed a total of 29,156 K-S tests between inter and intra-family distribution of similarity scores (7,289 families x [attention-based, CLS, mean, max] similarity metric]). Table 1 shows the summary statistics for the KS tests over all the protein families; the full table with individual family results can be found in the supplement. The HA vector method has the highest average and median value, showing that it is most distinguishable between in-family and random distributions. The p-values all show that these results are statistically significant given the number of tests conducted.

**Table 1:** We analyze the in-family vs. random family pairwise distances for all human protein families. This table reports the statistics of the KS tests between the in-family distribution and the random distribution, averaged over all human protein families. We see that the HA-based method has the highest mean KS test value, which shows that it has the highest separation between in-family and out-of-family.

|  | Min | Max | Mean | Median | Average p-value |
| --- | --- | --- | --- | --- | --- |
| Attention-based | 0.784 | 0.881 | 0.836 | 0.842 | $1.2 \times 10^{-6}$ |
| CLS | 0.700 | 0.820 | 0.760 | 0.770 | $1.6 \times 10^{-6}$ |
| Mean Pooled | 0.680 | 0.800 | 0.740 | 0.750 | $1.5 \times 10^{-6}$ |
| Max Pooled | 0.750 | 0.860 | 0.805 | 0.810 | $1.4 \times 10^{-6}$ |

### 5.4 Identifying HA sites can help predict active site location

We first evaluate the spatial distance between the HA site and the active site. Figure 8 shows the histogram of distances between HA sites and active sites for 5,594 proteins with an active site annotation in UniProt. For 85% of HA and active site pairs, they are at a euclidean distance of less than 12Å.

**Figure 8:**
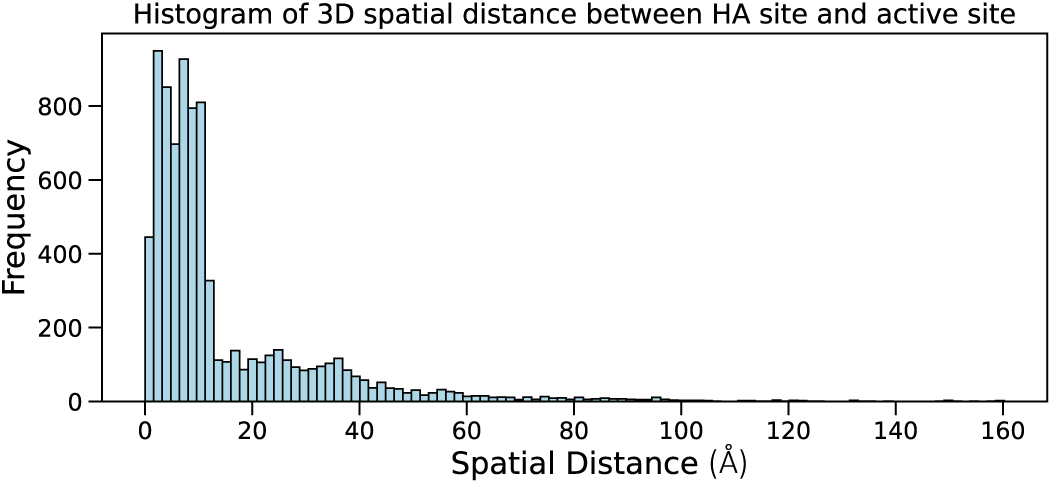
For all human proteins with active site annotation, we measure the spatial distance between the HA sites and their nearest active site and plot a histogram of these distances. 90% of HA site and active site pairs’ distance is less than 20Å and 85% of pairs are at a distance less than 12Å, showing that the HA sites and active sites are spatially close to each other.

Figure 9 shows three proteins with their HA sites annotated in red, active sites annotated in green, and any overlap is displayed in pink. These proteins have an HA site that overlaps with the active site, and the HA site residues are close to the active sites in the folded protein structure.

**Figure 9:**
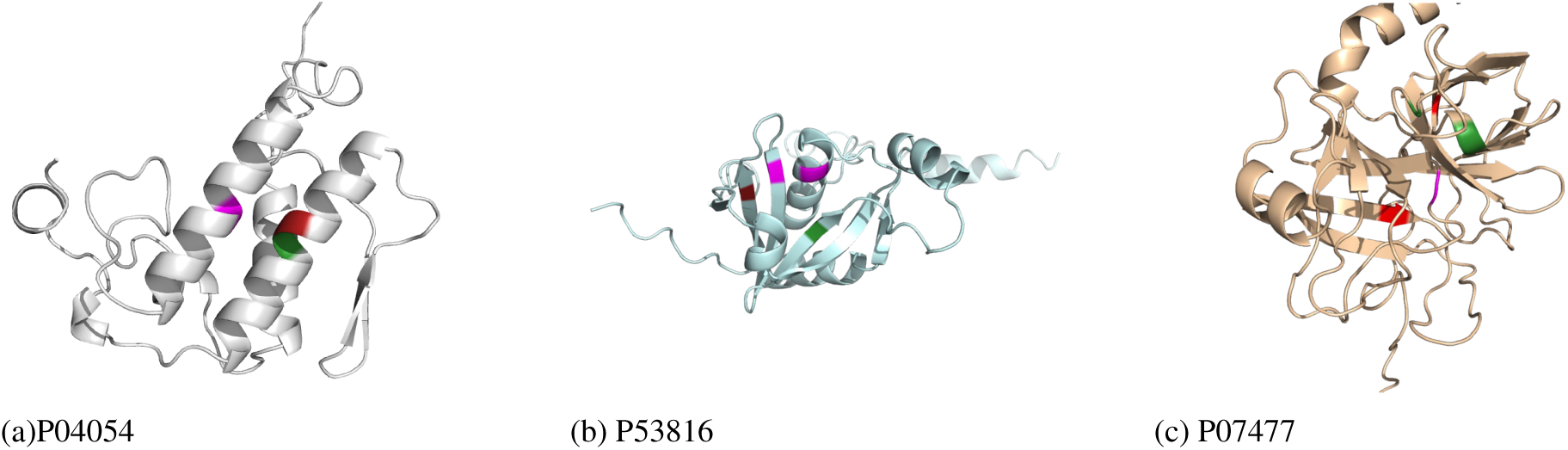
Three protein structures with active site annotations are visualized in 3-dimensional space. The HA sites are annotated in red, and the active sites are annotated in green. A residue that is both an HA site and an active site is annotated in pink. These structures show that the HA sites are in similar regions of the folded protein structure.

To show the utility of the HA sites, we use them to analyze two protein families that do not have an active site annotation in UniProt, methyltransferase (PF13847) and interferon-inducible GTPase (PF05049). Figure 10 shows the proteins in these two families, with their heatmaps and structures with HA sites annotated in red. In the methyltransferase family, we have two proteins of similar lengths (282 and 374) and one longer protein (689). Across all three protein examples, the layer of convergence identifies three HA sites on the beta strands. In QbN6R0, the longest example, the HA sites are separated by more AA than in the shorter examples, but the PLM identifies the same residues in all three examples. This family is a lysine methyltransferase and two of the three HA sites for all proteins are Lysine and S-Adenosyl methionine residues, which have been shown to be responsible for the catalytic function [34]. The specific residue annotations are not in UniProt, but upon investigation, they overlap with methionine and lysine that must be present in the catalytic pocket and require a nearby tyrosine, which is the third HA site in two of the protein [35]. For the GTPase family, the HA sites vary slightly between the two example proteins, as in one protein, two HA sites are in a beta-strand and alpha helix, while in the other protein, they are both on the beta-strand. In both proteins, the third HA site is within the domain region known the bind to magnesium, which can enhance GPT binding. All the predicted HA sites are within the G-domain, a conserved domain found in GTP-binding proteins [36, 37].

**Figure 10:**
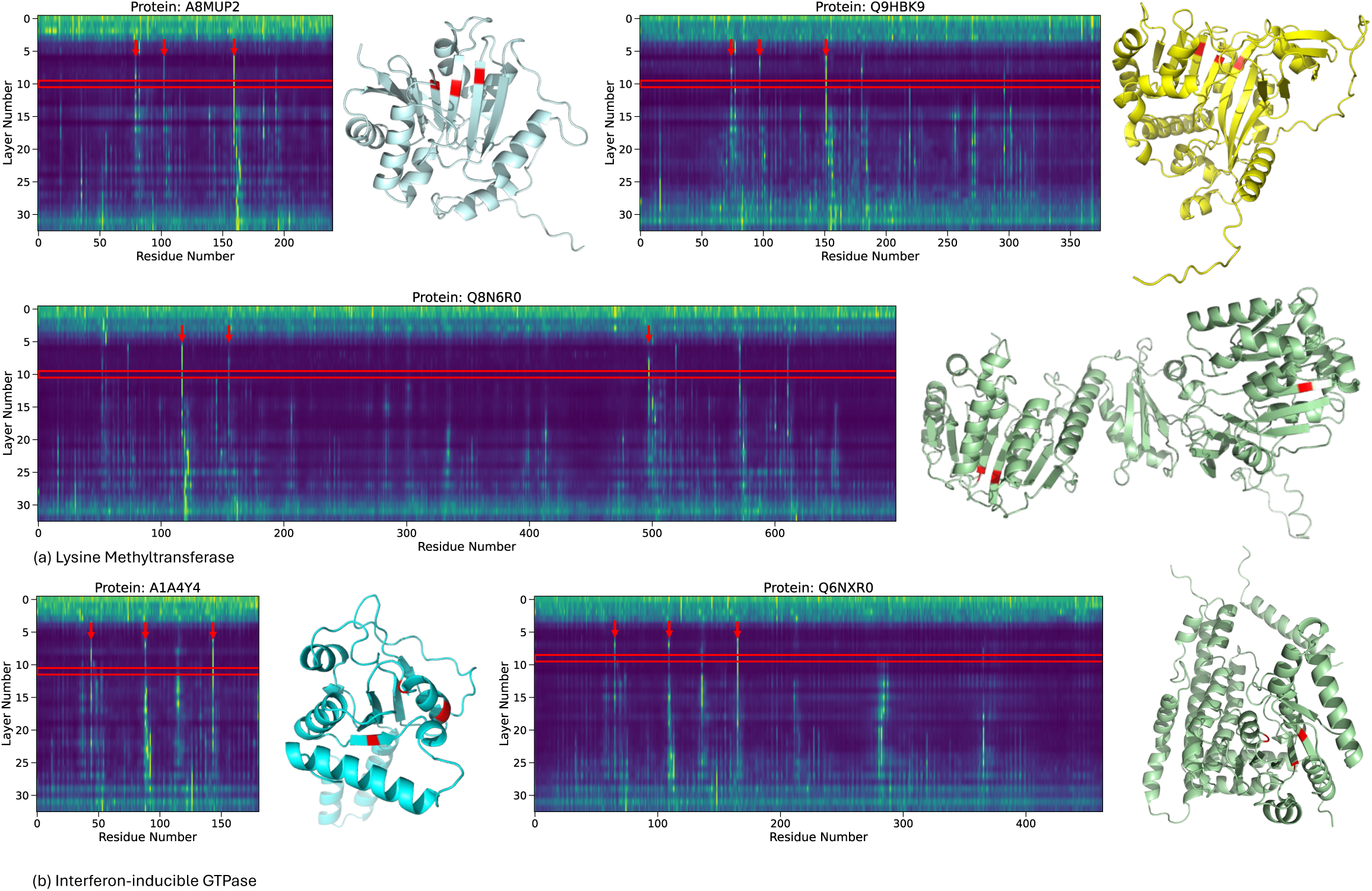
For two protein families (methylase and interferon-inducible GTPase) without active site annotations, we analyze the plausibility of the HA site as an active region of the protein. For each protein, we show the heatmap and the protein structure with the HA sites annotated in red. Panel A shows the methyltransferase family, and Panel B shows the GTPase family, with their heatmaps and structures annotated with HA sites in red. All proteins do not have an active site annotation in UniProt. For Panel A, the methyltransferase family, two shorter proteins (282 and 374 amino acids) and one longer protein (689 amino acids) share three HA sites on beta strands. Despite differences in sequence length, the same residues are identified by the PLM. Two HA sites correspond to lysine and S-Adenosyl methionine residues critical for catalysis [34], while the third site aligns with a tyrosine near the catalytic pocket [35]. In the GTPase family (Panel B), the HA sites differ slightly: one protein has two sites in a beta-strand and alpha helix, while the other has both on beta strands. A third HA site in both proteins is within the magnesium-binding region, enhancing GTP binding. All HA sites are located in the conserved G-domain [36, 37].

These results highlight the capability of HA sites to serve as reliable indicators of protein function. HA sites often correspond to residues directly involved in the protein’s activity, making them valuable targets for functional analysis.

### 5.5 Identification of HA sites precedes the PLMs identification of sequence and structure features typically associated with protein families

We find the association between the HA sites to the sequence motifs and structure of the protein; we compare at what attention layer the PLM attends to the HA sites versus the residues that are significant to the sequence and structure.

For the serine protease family (PF00089), we analyzed the multiple sequence alignments (MSAs), shown in Figure 11 with active sites annotated in green and high-attention (HA) sites annotated in red. The consensus of residues at each alignment position was displayed below the MSA, highlighting conserved and variable regions across the family. To enable detailed examination, specific sections of the MSA were zoomed in on and presented as Panels A, B, and C.

**Figure 11:**
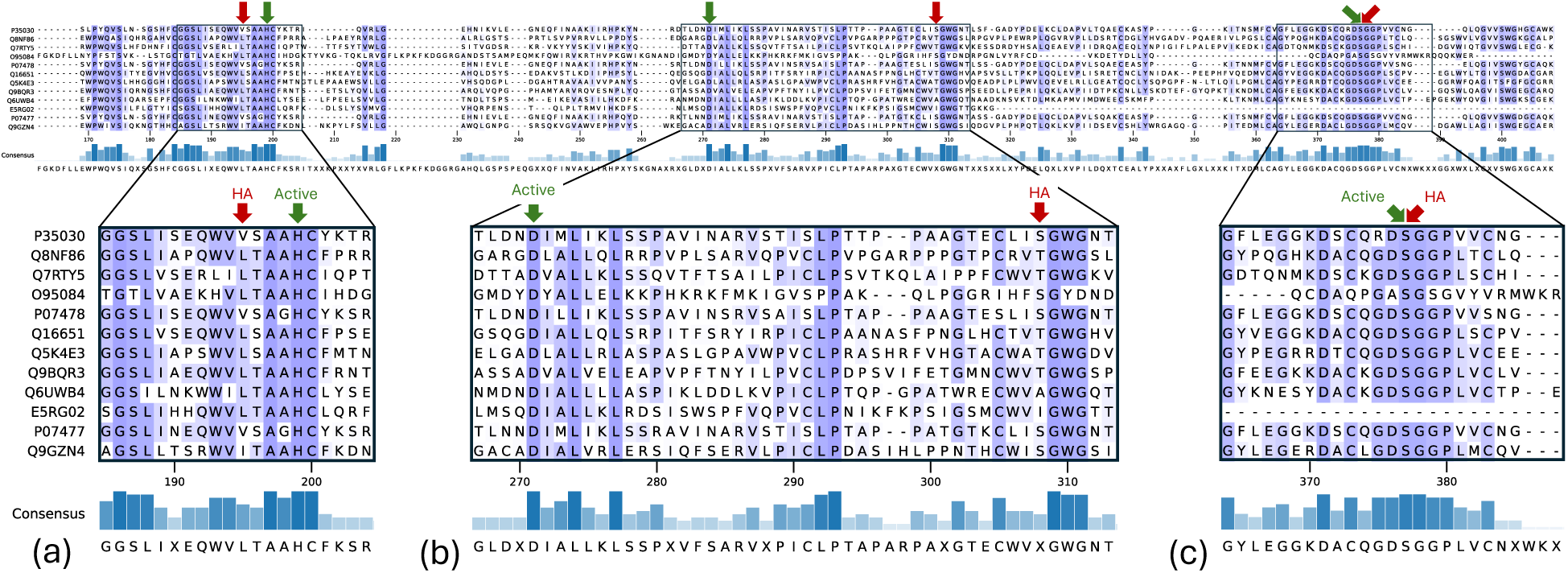
For the serine protease family (PF00089), we show the full multiple sequence alignments with active sites annotated in green and HA sites annotated in red. The consensus of residues at each site is shown below the alignment. Panels A, B, and C show a zoomed-in section of the MSA to facilitate visualization of the percent consensus at each HA site and the relative position between the HA sites and active sites. In panel A, the percent consensus of the HA site is not the highest within that region and the HA site is within 4 residues of the active site. In panel B, the HA site is again at a low consensus site but is further from the nearest active site. However, this HA site occurs right before the *GWG* residue pattern that is often found within the active site of serine proteases, as the Glycines play a role in the structure of the pocket and the Tryptophan within the pocket interacts with hydrophobic amino acids on the substrate and influences the specificity [38, 39, 40]. Panel C is on the active site, which shows the functional importance of this HA site. These panels emphasize the percent consensus at each HA site and their spatial relationship to active sites.

In Panel A, the HA site exhibits moderate percent consensus and is located within four residues of the nearest active site. Panel B highlights an HA site with lower consensus that is positioned further from the nearest active site. Interestingly, this HA site occurs just before the conserved GWG motif, a sequence pattern frequently found within the active pocket of serine proteases [38, 39, 40]. The glycines in this motif are responsible for the shape of the catalytic pocket, while the tryptophan interacts with hydrophobic residues on the substrate, influencing the specificity of substrate binding. Panel C presents an HA site that overlaps exactly with an active site. This shows that the HA sites are not solely governed by the regions of high consensus, and thus PLM attends to features outside of simple sequence similarity early on in the model.

Figure 12 panel A shows a histogram of the consensus at each HA site for all protein families. 90% of all HA sites have a 50% or lower consensus. For the HA sites that are at 100% residue consensus, 92.7% are overlapping with a known active site. A high rate of consensus is expected for active sites, as they are typically conserved across all proteins in the family.

**Figure 12:**
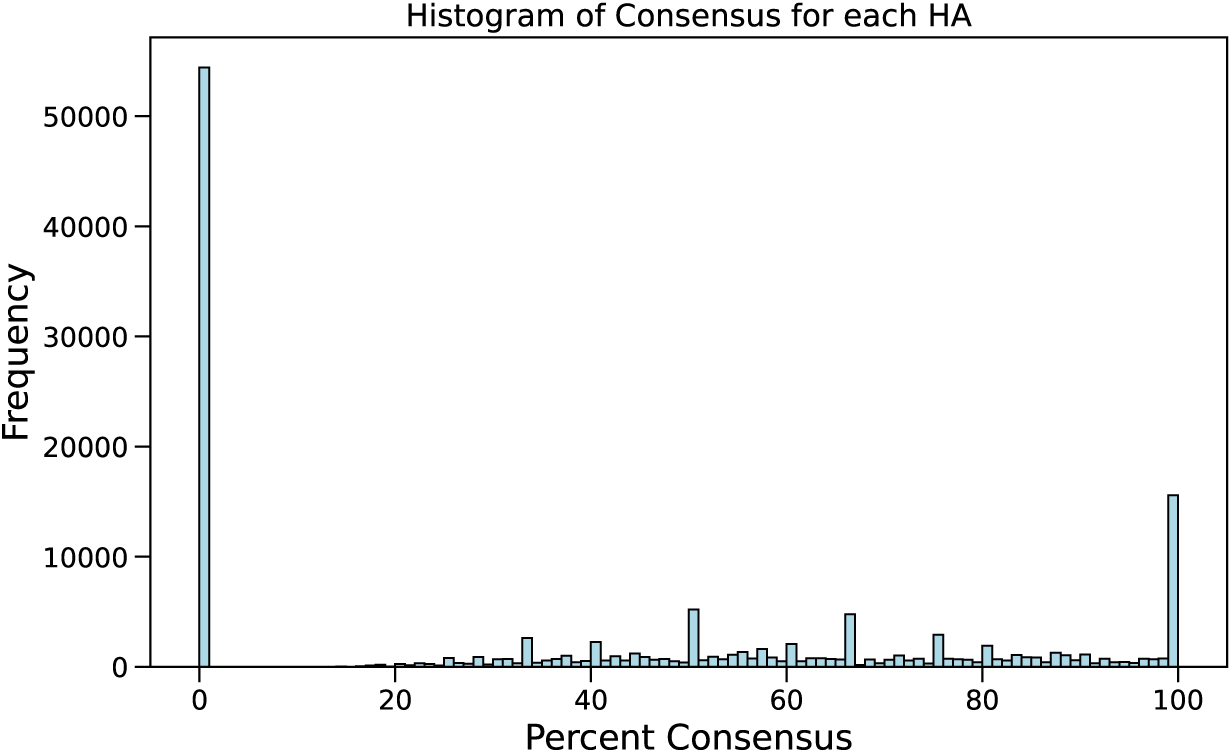
Histogram of the percent consensus among sequences in the family of the residue at each HA site for all proteins in the human proteome. The majority of HA sites occur close to a 0 percent consensus, showing that the HA sites are not dominated by the conserved sequence regions. The next most frequent percent consensus is 100%, and 93% of these residues are also known active sites (from UniProt). These results suggest that HA sites are not strictly determined by highly conserved residues but are enriched for active residues when conservation does occur.

To compare the HA sites to the structure of the protein, we correlate the attention matrix values to the contact map. Figure 13 A shows the correlation between the attention matrix and the contact map over each layer, where each line indicates the correlation values for an individual protein and the black line is the average across all proteins. The correlation tends to oscillate around constant value for most of the network layers and there is a pronounced uptick in correlation at the last layer. Furthermore, Figure 13 B shows a histogram of the layer with the highest correlation value between the attention matrix and the contact map, and 94.7% of proteins have the highest correlation at the last layer. Because the ESM-2 model uses regression on the attention matrices to generate the contact map, we do not expect the attention matrices to be exactly equivalent to the contact map, explaining the lower levels of correlation. However, the increase in correlation over the layers shows that the early attention layers are not significantly correlated to the structure in comparison with the last layer of attention. In the range of the Layers of Convergence across all protein families (Layer 9-10) the average correlation values are between 0.280 and 0.321, with p-value less than 2.12 × 10*^−^*^4^. These findings suggest that the HA sites and the PLM’s ability to detect them in the early layers of the model play a crucial role in the model’s decision-making process and indicate the HA sites’ relevance to the protein’s function.

**Figure 13:**
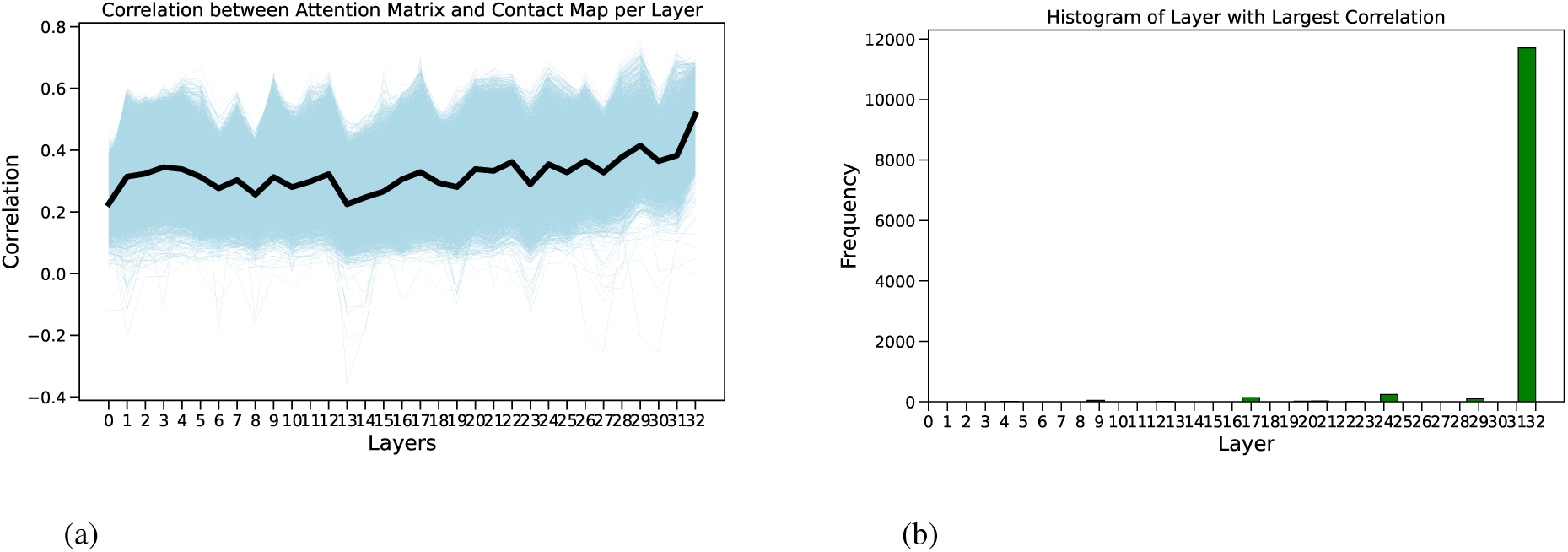
Panel A shows the correlation between the attention matrix and the contact map over each layer for all human proteins. The blue plot lines are the correlate on of the attention matrix and the contact map for each protein, and the black line is the average over all proteins. There is no significant correlation through the layer, but at layer 33, the average correlation increases significantly. This shows that the last layer is the most related to the structure, rather than the middle layer, which is where the layer of convergence occurs. To further elucidate this, Panel B shows a histogram of the layer with the largest correlation between the attention matrix and the contact map for all human proteins. The majority of proteins have the highest correlation at the last layer. Therefore, the convergence layer that occurs in the middle layers of the model is not necessarily correlated to the structure of the protein, and so the HA sites are not necessarily structurally important residues.

## 6 Discussion

We conclude that HA sites are ubiquitous across all human proteins, and the PLM relies on their identification to categorize the protein into groups. HA sites are (1) found across all proteins in the human proteome, (2) reliably near important functional regions of the protein, and (3) robust across protein families with a wide range of sequence lengths. The PLM converges on the HA sites early on in the attention layers and then generates more fine-grained attention to distinguish specific changes in the sequence. These HA sites are reliably consistent across members of the same protein family. We provide the HA sites for all proteins and the protein family alignments for the entire human proteome, which can be found here: https://github.com/Helix-Research-Lab/ESM_highAttention.

A common task with PLMs is to use the resulting vectors to find similarities between proteins. Using the normalized attention at the HA sites creates a tighter cluster for in-family proteins than using the representation vectors. This is because the representation vectors are a result of the last layer of attention, which is locally specific rather than globally specific. Therefore, the representation vectors are more suited to study the local structure of the protein, than the global functional family of the protein. To correctly distinguish functional similarities between proteins, the HA sites should be used.

The HA sites are also correlated in 3D space to the active sites of the protein. This follows from the notion that the HA sites are efficient protein family identifiers, as the active sites are essential to the function. We propose that because the HA sites are or are near known active sites, they can be used to identify regions of the protein that are potential active sites. We have shown the functional relevance of the HA sites for two protein families with no active site annotation and shown that the HA sites identify the residues that perform similar functions regardless of the length of the proteins.

The attention layers at the end of the model are correlated to the structure of the protein, and thus they contain the specific local pairwise relationships that are important to the protein folds. Instead, the HA sites emerge in the early attention layers as clear signals, with all other pairwise attention at very low values. These HA sites are not correlated to the location of high sequence conservation. We have shown that the HA sites can be used to define the protein family but are not correlated with statistical significance to the structure or the sequence definition of the protein family. Thus, we conclude that the HA sites are early detectors of the function of the protein.

Because the ESM model is trained on all protein sequences, the model has seen examples of all protein families, whether or not they are known as a functionally similar set. Therefore, the model is able to detect the sequence relationships and latent features that characterize each family. In inference, when a protein is introduced to the model, the first features to emerge are those that classify the protein into its broader family, which occurs at the layer of convergence, and then more detailed features emerge as the model progresses.

While the described method and analysis make gains in explaining the PLM and identifying functional residues, we recognize opportunities for improvement. Our analysis is limited to the first layer, the layer of convergence, with high attention values for functionally relevant residues, but is limited in discussion of the progression of attention as the model progresses past the layer of convergence. The secondary attention sites that emerge are likely to be informative of functional residues that are more specific and based on the local context of the protein. Furthermore, our analysis focuses on the ESM model, as it offers access to the contact maps. While we expect other bidirectional models (BERT-style) to function similarly, the specific parameters of the model could influence which features have high attention. To this end, we use ProtTrans, a BERT-style PLM [41], to get the attention matrices for an example protein family, PF00089, and we find that for the trypsin-like serine protease family, the identified HA-site residues are the same. However, this analysis can be expanded upon to evaluate any changes across all human proteins. Furthermore, we evaluate only one type of ESM model without considering other specifications of parameters (number of layers, number of parameters, dimension size). These hyperparameters of the trained model may influence the specificity of the high-attention sites. We expect that a larger model would identify similar features in the layer of convergence as they remain the features that define the family, but that the layer of convergence would be achieved earlier and subsequent high-attention sites would represent more detailed, local interactions. This method provides a necessary first step in explainability for PLMs by evaluating the attention progression and identifying a biological reason for the early high-attention sites across the entire human proteome.

## Acknowledgments

We performed the data download and generation, method development, and analysis on the Sherlock cluster, and we thank Stanford University and Stanford Research Computing Center for the computational resources.

## 7 Data Availability

There is no primary data used in this study. All data and code are made available at https://github.com/Helix-Research-Lab/ESM_highAttention.

